# Consequences of decoy site repair and feedback regulation on neurotransmission dynamics

**DOI:** 10.1101/2024.07.20.604401

**Authors:** Oliver Gambrell, Abhyudai Singh

## Abstract

Neurons form the fundamental unit of the central nervous system with the human brain containing close to 100 billion neurons. We present a systems-level model of a chemical synapse by which signals from a presynaptic neuron are transmitted to a postsynaptic neuron. In this model, neurotransmitter-filled synaptic vesicles (SVs) dock with a given rate at a fixed number of docking sites in the axon terminal of the presynaptic neuron. Upon the arrival of an action potential (AP), each docked SV has a certain probability to fuse with the presynaptic membrane and release neurotransmitters into the synaptic cleft. After the SV fusion event, the corresponding docking site undergoes repair before becoming available to be reoccupied by an SV. We develop a stochastic model of these coupled processes and derive exact analytical results quantifying the mean and the degree of random fluctuations (i.e., noise) in the levels of docked SVs and released neurotransmitters in response to a train of APs. Our results show that the repair of docking sites exacerbates synaptic depression, i.e., reduces the ability of the chemical synapse to release neurotransmitters in response to an AP. Moreover, repair amplifies statistical fluctuations in neurotransmission for fixed mean neurotransmitter levels. We next consider feedback regulation where the released neurotransmitters affect the rate of SV docking. Counterintuitively, our analysis reveals that for certain physiological parameter spaces, positive feedback loops can reduce noise levels in both the number of docked SVs and neurotransmitters in the cleft.

## I. Introduction

Chemical synapses form the fundamental basis by which neurons communicate. This communication occurs through neurotransmitter-filled synaptic vesicles (SVs) present at the axon terminal of the presynaptic neuron. These SVs fuse with the presynaptic axon membrane to release their neurotransmitter content into the synaptic cleft (Fig. 1). This release could occur spontaneously or in response to an action potential (AP), and the released neurotransmitter impacts the activity of the postsynaptic neuron, for example, by opening ion channels in the target cell.

**Fig. 1.**
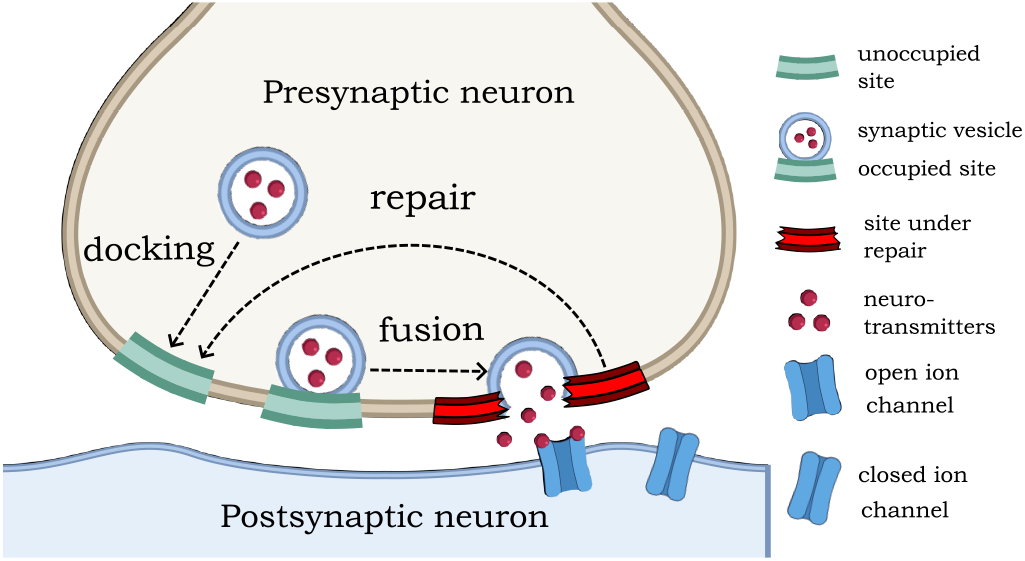
Model schematics of a chemical synapse with docking site repair. Synaptic vesicles (SVs) are shown to dock at unoccupied (empty) docking sites in the axon terminal of the presynaptic neuron. The arrival of an AP triggers SV fusion to the membrane and the release of its neurotransmitter cargo into the synaptic cleft impacting the activity of the postsynaptic neuron. After SV fusion, the docking site undergoes repair with a given rate before again becoming available to be reoccupied by SVs.

The simplest model capturing the dynamics of neurotransmitter release consists of SVs continuously docking at a fixed number of docking sites in the axon terminal, and the arrival of an AP triggers SV fusion and neurotransmitter release. In response to a train of APs, the number of docked SVs decreases (a phenomenon termed as synaptic depression) and reaches a dynamic equilibrium set by the balance of AP-triggered depletion and SV replenishment [1]–[4]. Prior work has analyzed a stochastic version of this model considering stochasticity in AP arrival times, probabilistic SV docking/fusion, and systematically investigated how different model parameters impact the degree of statistical fluctuations in SV counts [5]–[8]. Furthermore, the impact of stochastic neurotransmitter releases on the timing precision of AP generation in the postsynaptic neuron [9]–[11] was investigated using standard integrate-and-fire models [12]–[14].

It is important to point out that these works assume a single SV pool, where each docked SV acts identically, and independently of each other. However in reality, there could be multi-types of SV pools and significant site-to-site differences in SV docking and fusion properties [15]– [24] that play key roles in defining the neurotransmission dynamics at different frequencies of presynaptic APs [25]– [30].

One limitation of our previous work was that once a docked SV fuses, that docking site becomes immediately available to be re-occupied by a SV. In this work, we introduced a *docking site repair state*, where after SV fusion, the site is unavailable for SV docking. This biologically corresponds to the reassembly of the molecular machinery needed for SV docking and fusion [31] and leads to the model shown in Fig. 1, where a SV fusion event results in the site immediately transitioning to a repair state. Sites are assumed to be repaired with a given *repair rate*, after which they transition into an empty state, and SV docking at empty sites occurs with a given *refilling rate* or *replenishment rate*. This results in a four-dimensional stochastic system with state variables being the number of docked SVs (i.e., occupied docking sites), the number of sites under repair, the number of empty sites, and the number of neurotransmitters in the cleft. Since the total number of docking sites is conserved and assumed to be fixed, this model can be reduced to a three-dimensional system. Stochastic analysis of this system using both analytical approaches and numerical simulations shows that the repair state can enhance statistical fluctuations in the number of docked SVs and released neurotransmitter counts for fixed mean levels. This amplification is only seen for a high number of docking sites and occurs at an optimal value of the repair rate.

We further generalize the model to consider feedback regulation in the SV docking rate via neurotransmitters. For example, high levels of neurotransmitter buildup in the synaptic cleft reduce the SV docking rate, thus limiting the number of SV fusing for the next AP. Our analysis shows that positive feedback is always better than negative feedback in minimizing fluctuations in the number of docked SVs. In contrast, depending on the relative time scales of AP arrival frequency and the neurotransmitter decay rate, either feedback type (positive or negative) can function to buffer statistical fluctuations in the neurotransmitter levels.

### II. Mechanistic modeling OF Chemical synapse

The basic model formulation of a chemical synapse consists of a given number of docking sites *M* ∈ {1, 2, …} in the active zone of the presynaptic axon terminal. Each site can be in one of three possible states (Fig. 1):

- Site is *occupied* by a docked SV.
- Site is under *repair* and unavailable for SV docking.
- Site is *empty* and ready for SV docking.

We let integer-valued random processes *n*(*t*) and *r*(*t*) denote the number of docking sites that are occupied and undergoing repair at time *t*, respectively. Given that the total number of sites is fixed, the number of empty sites is *M* − *n*(*t*) − *r*(*t*). The number of neurotransmitters in the synaptic cleft at time *t* is denoted by *z*(*t*).

The overall model consists of four events that occur randomly over time as prescribed by their stochastic propensities defining the frequency with which they occur. When an event occurs the state-space {*n, r, z*} undergoes an integer-valued jump or reset. These events and corresponding resets are as follows:

### 1) AP arrival

APs are assumed to arrive as per a Poisson process with frequency *f*, i.e., the inter-AP arrival times are exponentially distributed with mean 1*/f*. AP arrival triggers each docked SV to fuse and release neurotransmitters with *release probability p*_*r*_. Given *n* docked SVs, each with SV fusing probability *p*_*r*_, the number of SVs fusing is a binomially distributed random variable

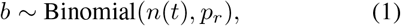

where *b* ∈ {0, 1, …, *M*} is often referred to in literature as the quantal content. Conditioned on *n*(*t*), the probability mass function (pmf) of *b* is

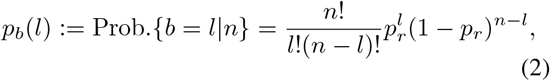

*l* ∈ {0, 1, …, *n*}. This event results in the following reset

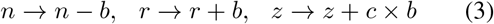

corresponding to docked SV depletion by *b*, an exact similar increase in sites undergoing repair, and an increase in neurotransmitter levels in the cleft, respectively. Here *c* is the number of neurotransmitters per SV and is assumed to be a fixed positive integer.

### 2) SV docking

Each empty docking site is replenished by SVs with a *refilling rate k*. Recalling that *M* − *n* − *r* is the number of empty docking sites, this event occurs with a propensity *k*(*M* − *n* − *r*), i.e. the probability that it will occur in the next infinitesimal small interval (*t, t* + *dt*] is *k*(*M* − *n* − *r*)*dt* and the event results in the reset

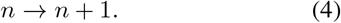

This formulation implies that the time taken by an empty site to become SV occupied is exponentially distributed with mean 1*/k*.

### 3) Site repair

Once a docked SV fuses, it’s corresponding site undergoes repair with *repair rate γ*_*r*_. In the stochastic model formulation, repair events occur with propensity *γ*_*r*_*r*(*t*), and upon repair completion *r*(*t*) is reset as

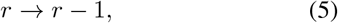

implying each site’s repair time is exponentially distributed with mean 1*/γ*_*r*_.

### 4) Neurotransmitter clearance

Finally we assume that each neurotransmitter in the cleft is degraded (cleared) with rate *γ*_*z*_. This event occurs with propensity *γ*_*z*_*z*(*t*) and corresponding reset

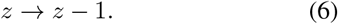

Fig. 2 illustrates the time evolution of the stochastic processes *n*(*t*), *r*(*t*), *z*(*t*) as per the above model for given parameter values. Note that between two successive APs, the number of sites under repair *r*(*t*) monotonically decreases with time, while the number of occupied sites *n*(*t*) monotonically increases. For the reader’s convenience, state variables and model parameters are summarized in Table I.

**Fig. 2.**
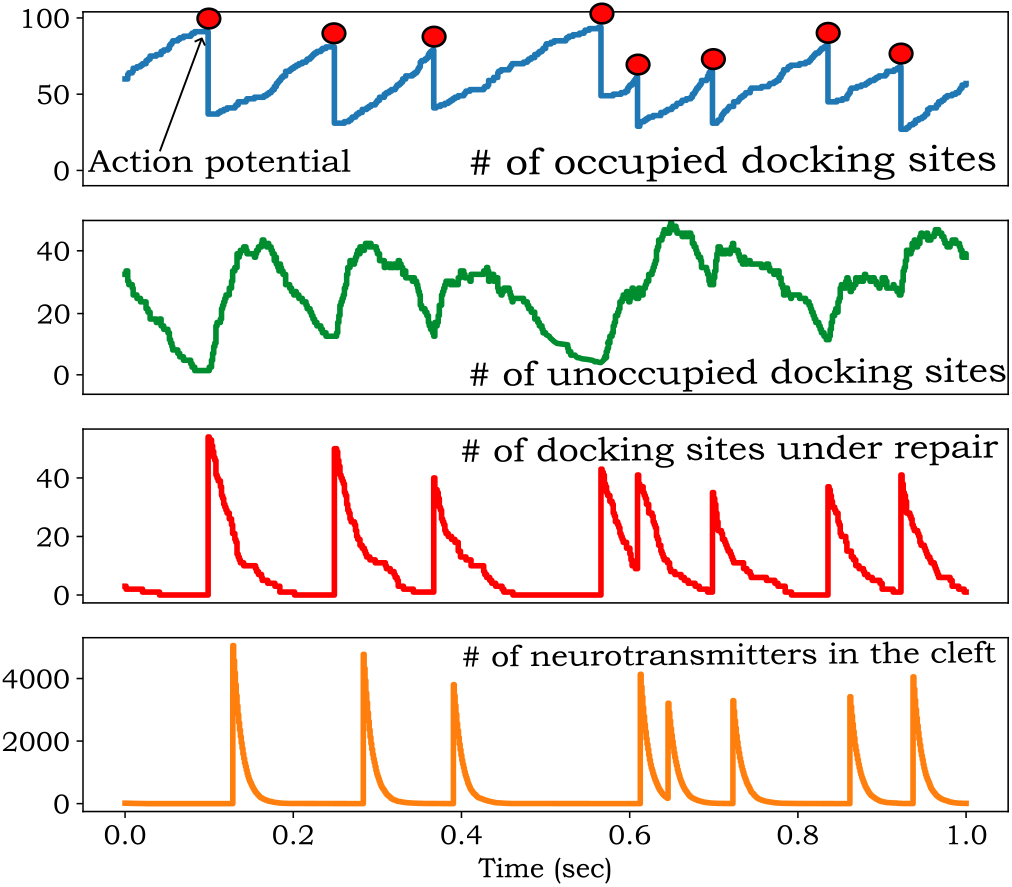
Sample realization of the random processes underlying the stochastic model of a chemical synapse. In response to a train of presynaptic APs (red dots on the top plot), docked SVs fuse into the axon membrane releasing their neurotransmitter content into the synaptic cleft. Plots from top to bottom represent the number of SV-occupied docking sites, empty sites (available for SV docking), sites under repair (unavailable for SV docking), and the number of neurotransmitters in the cleft, respectively. The parameters used for this simulation are: *M* = 100, *k* = 10 sec^*−*1^, *γ*_*r*_ = 40 sec^*−*1^, *p*_*r*_ = 0.5, *γ*_*z*_ = 100 sec^*−*1^, *c* = 100, *f* = 10 *Hz*.

**TABLE 1.** Parameters used in the stochastic model of a chemical synapse.

| Parameter | Description |
| --- | --- |
| $M$ | Number of docking sites for synaptic vesicles (SVs) in the axon terminal. |
| $f$ | Poisson rate of action potential (AP) arrival at the active zone of the axon terminal. |
| $n(t)$ | The number of docked SVs at time $t$. |
| $r(t)$ | The number of docked sites undergoing repair and not available for SV docking at time $t$. |
| $p_r$ | AP-triggered probability of docked SV fusion resulting in the release of its neurotransmitter cargo. |
| $z(t)$ | The number of neurotransmitters in the synaptic cleft at time $t$. |
| $k$ | The refilling rate of SV docking at empty docking sites. |
| $c$ | The number of neurotransmitters per SV. |
| $\gamma_r$ | The repair rate at which docking sites are repaired. |
| $\gamma_z$ | The rate of neurotransmitter clearance in the synaptic cleft. |
| $\langle \rangle$ | Expected value of random variables and random processes. The notation $\overline{\langle \rangle}$ is used for steady-state expected values. |
| $FF_n, FF_z$ | The steady-state Fano factor (variance divided by the mean) of $n(t)$ and $z(t)$, respectively. |

## III. Quantifying statistical fluctuation in population counts

Having defined the model describing the time evolution of coupled integer-valued random processes *n*(*t*), *r*(*t*), *z*(*t*), we are primarily interested in quantifying their statistical moments, particularly focusing on the first- and second-order moments only. Using the angular brackets ⟨ ⟩ to denote the expected value operation, ⟨*n*(*t*)⟩, ⟨*r*(*t*)⟩, ⟨*z*(*t*)⟩ represents the transient mean population counts. The average population counts at steady-state are denoted by

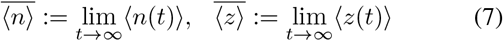

with 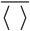 denoting the moments in the limit *t* → ∞ whenever the limit exists. We quantify the magnitude of statistical fluctuations using the steady-state Fano factors (variance divided by the mean) defined as

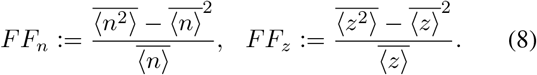

Given a set of events with occurrence propensities and corresponding jumps in the state space, the time evolution of statistical moments can be straightforwardly derived using the tools of moment dynamics. Referring the reader to [32] for more details, for given integers *i*_1_, *i*_2_, *i*_3_ ∈ {0, 1, 2, …}, the time derivative of the uncentered moment 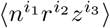 is

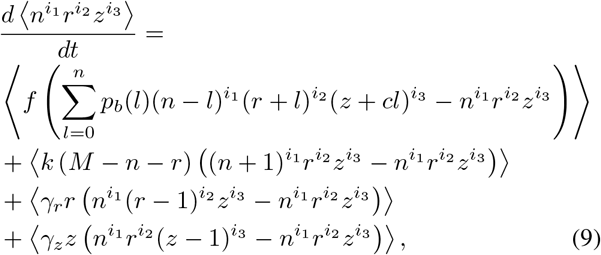

which is in essence the expected value of the change in the monomial 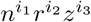 when an event occurs multiplied by its propensity, summed up across all the four classes of events defining the stochastic model.

## IV. Average population counts

The time evolution of the mean levels can be derived by appropriately selecting integers *i*_1_, *i*_2_, *i*_3_. For example, taking *i*_1_ = 1, *i*_2_ = 0, *i*_3_ = 0 in (9) yields

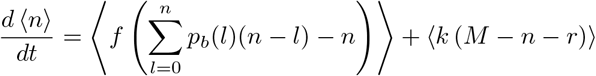

which using the pmf (2) simplifies to

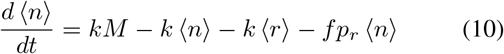

given that

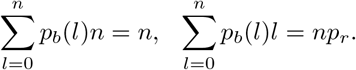

Similarly, one can derive

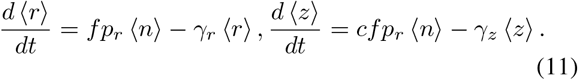

The solution to the mean dynamics (10)-(11) is shown in Fig. 3 for increasing docking site repair time. Solving (10)-(11) at steady state yields the following steady-state mean levels

**Fig. 3.**
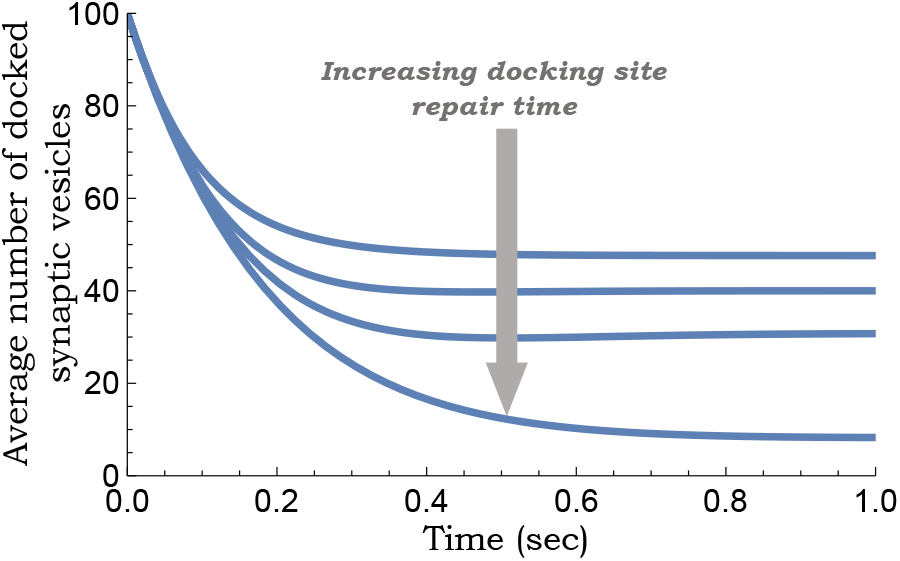
Repair of docking sites exacerbates synaptic depression Statistics. The average number of docked SVs ⟨*n*(*t*)⟩ as obtained by solving (10)-(11) for increasing docking site repair time 1*/γ*_*r*_. Parameters taken as *M* = 100, *p*_*r*_ = 0.5, *f* = 10 *Hz, k* = 5 *sec*^*−*1^ and *k*_*r*_ = 50, 10, 4, 0.5 *sec*^*−*1^ from top to bottom.

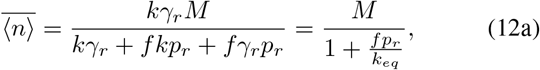

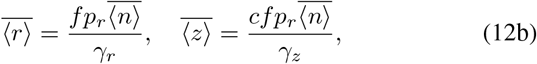

which can be rewritten in terms of an equivalent rate *k*_*eq*_, defined as

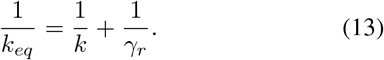

Note from (12a) that the balance between the rate of SV deletion (*fp*_*r*_) and the equivalent rate of site refilling (*k*_*eq*_) determines the docked SV count at equilibrium.

## V. Statistical fluctuations in population counts

We next focus our efforts on quantifying statistical fluctuations by writing the moment dynamics of second-order moments. For example, taking *i*_1_ = 2, *i*_2_ = 0, *i*_3_ = 0 in (9) yields

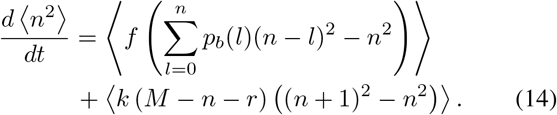

Using the fact that

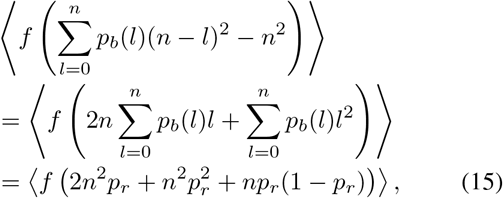

(14) can be written as

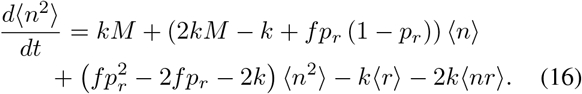

The standard way to solve these moment dynamics is to define a vector *µ* that contains all first- and second-order moments

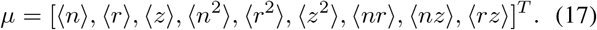

Then, the time evolution of *µ* is given by a linear dynamical system

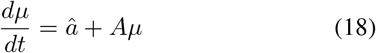

that can be solved both transiently and at steady-state [33], [34]. Performing the steady-state analysis in *Wolfram Mathematica* yields an exact analytical expression for the Fano factors (8), but these expressions are too complicated to be presented here. The special case of *γ*_*r*_ → ∞ (i.e., ignoring docking site repair or instantaneous site repair) was analyzed previously in [8] and the steady-state Fano factor of docked SVs was reported as

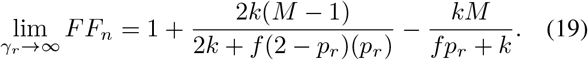

We refer the reader to [8] for a systematic analysis of *FF*_*n*_, where *FF*_*n*_ is a function of model parameters when docking site repair is ignored.

Our goal is to understand how repair processes (i.e., finite *γ*_*r*_) impact the Fano factors. Interestingly, our analysis shows that when *M* = 1 the Fano factor for docked SVs as obtained by solving (18) is given by

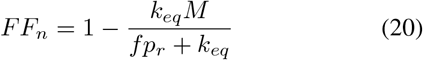

and only depends on the site repair rate *γ*_*r*_ through *k*_*eq*_. Note that (20) is simply (19) with *k* replaced by *k*_*eq*_ for *M* = 1. However, this property is restricted to *M* = 1 and is not true for larger values of *M*. In Fig. 4 we plot the steady-state Fano factors of *n*(*t*) and *z*(*t*) as a function of *γ*_*r*_ for *M* ≫ 1. In this plot, as *γ*_*r*_ is varied, the equivalent rate *k*_*eq*_ is held fixed by simultaneously changing *k* as per (13) ensuring fixed mean levels 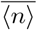 and 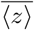. Our results show:

**Fig. 4.**
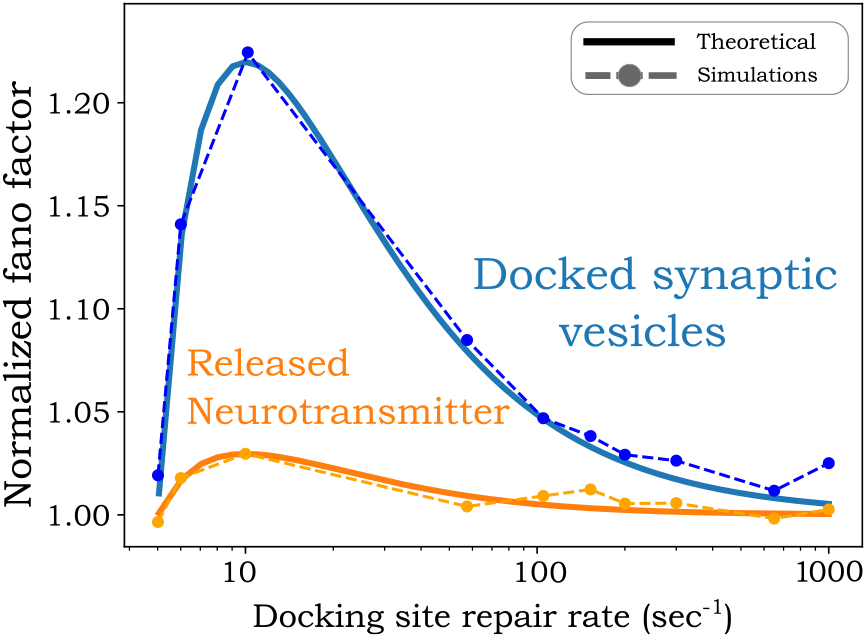
Docking site repair enhances stochastic fluctuations in population counts for fixed mean levels. The plots of the steady-state Fano factors *FF*_*n*_ and *FF*_*z*_ as a function of the docking site repair rate *γ*_*r*_. The Fano factors were obtained from solving the moment dynamics (18). As *γ*_*r*_ varies, SV docking rate *k* is also varied to keep *k*_*eq*_ in (13) fixed. The Fano factors are normalized by their values for instantaneous site repair (i.e., *γ*_*r*_ → ∞). Other parameters are taken as *M* = 100, *p*_*r*_ = 0.5, *f* = 10 *Hz, k*_*eq*_ = 5 *sec*^*−*1^, *c* = 100 and *γ*_*z*_ = 100 *sec*^*−*1^. The corresponding normalized Fano factors obtained from running 10^3^ sample realizations of the stochastic model are shown as dots on top of the analytical results.

- Statistical fluctuations in both docked SV and neurotransmitter levels can vary non-monotonically as a function of *γ*_*r*_ (Fig. 4).
- The steady-state Fano factors are maximized when *k*_*r*_ = 2*k*_*eq*_.
- In the limit *M* → ∞, *γ*_*z*_ → ∞, the values of *FF*_*n*_ and *FF*_*z*_ when *k*_*r*_ = 2*k*_*eq*_ (normalized by their values for instantaneous site repair) are as follows

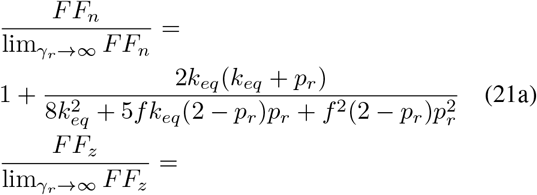

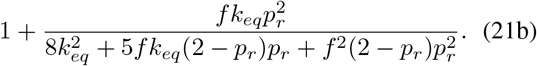

Thus, (21) represent the maximum amplification in statistical fluctuations as a result of docking site repair that occurs at *k*_*r*_ = 2*k*_*eq*_.

## VI. Feedback regulation of neurotransmission

Having quantified the impact of docking site repair on neurotransmission fluctuations we now consider feedback regulation. Recall the refilling rate *k* with which SVs dock at empty sites. To incorporate the feedback we modify this rate to be a function *k*(*z*) of the neurotransmitter level in the synaptic cleft. We consider *k*(*z*) to be a monotonically increasing or decreasing function of *z*(*t*) that implements positive or negative feedback, respectively.

Given the nonlinearities introduced via feedback, we resort to approximations, such as the Linear Noise Approximation (LNA) [35]–[37] to obtain moments. Considering small fluctuations in state variables, one can linearize the docking rate around the steady state mean

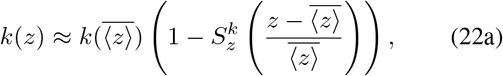

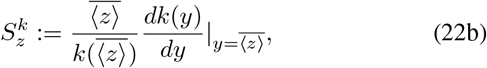

where 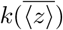 is docking rate at the mean neurotransmitter level, and 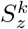 is the log sensitivity of *k*(*z*) with respect to *z* evaluated at its mean and interpreted as the *feedback strength*. Positive and negative values of 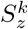 correspond to positive and negative feedback, respectively, with 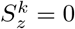 recovering the no feedback case. The LNA involves linearizing the propensities around the steady-state mean level and then using the linearized propensities to derive and solve the resulting moment dynamics [35].

To investigate the impact of feedback we consider the Fano factors normalized by their corresponding value in the absence of feedback

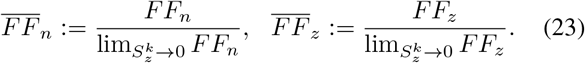

The derivatives

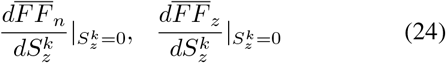

of these normalized Fano factors with respect to 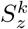 evaluated at 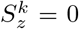 is plotted in Fig. 5 as a function of two dimensionless quantities: *f/γ*_*z*_ (the relative rate of AP arrival with respect to the neurotransmitter decay rate) and 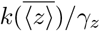 (the relative rate of SV docking with respect to neurotransmitter decay rate). In this plot, positive (negative) values of the derivatives (24) imply noise buffering is mediated by negative (positive) feedback.

**Fig. 5.**
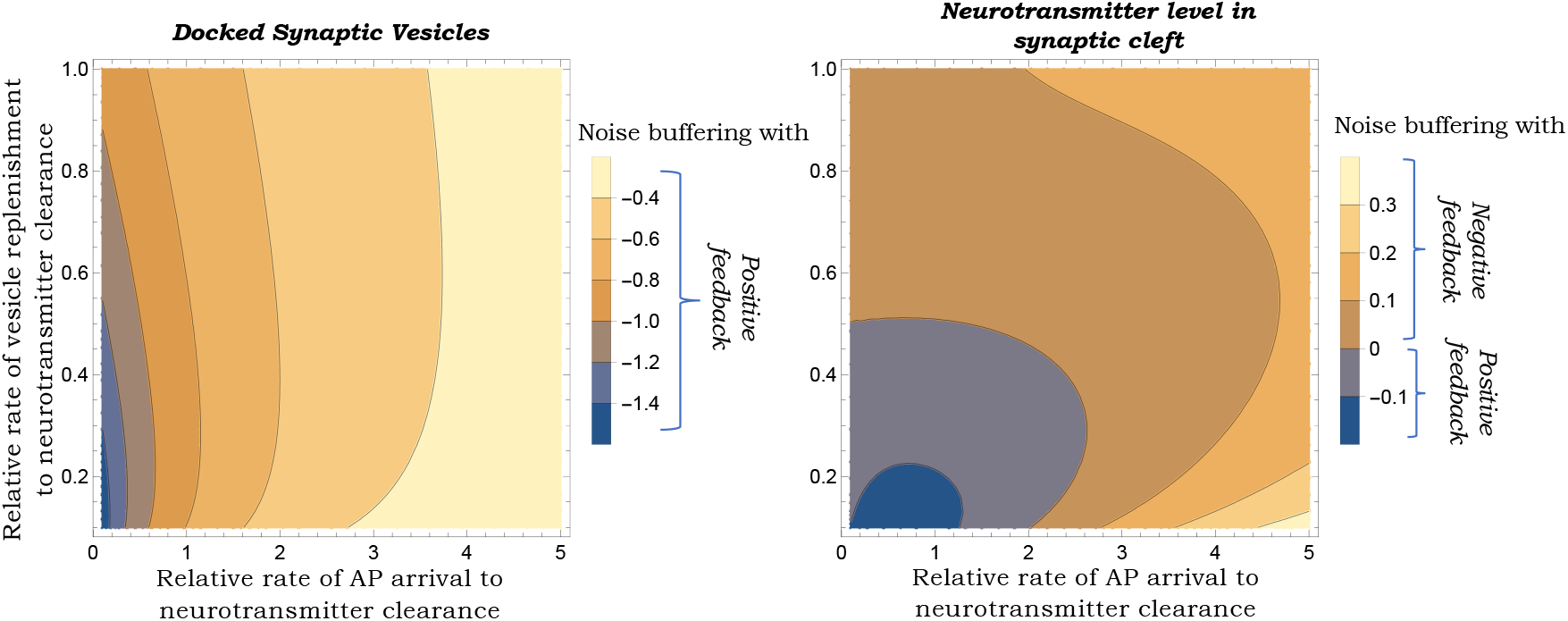
Positive feedback regulation in neurotransmission can attenuate statistical fluctuations in docked SV counts. Plots of the derivatives (24) of the docked SV Fano factor (left) and neurotransmitter level Fano factor (right) as a function of dimensionless quantities *f/γ*_*z*_ (the relative rate of AP arrival with respect to the neurotransmitter decay/clearance rate) and 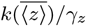 (the relative rate of average SV refilling rate with respect to neurotransmitter decay/clearance rate). The steady-state Fano factors were obtained using the LNA that involves linearizing the event propensities considering small fluctuations around steady-state mean levels. In this case, positive feedback always reduces fluctuations in docked SV counts (left) but either negative/positive feedback can reduce fluctuations in neurotransmitter levels in the cleft (right). For this plot we assume instantaneous site repair *γ*_*r*_ → ∞ and parameters *M* = 100, *p*_*r*_ = 0.5, *c* = 100 and *γ*_*r*_ = 100 *sec*^*−*1^.

In the limits *M* → ∞, *γ*_*z*_ → ∞, *γ*_*r*_ → ∞ (i.e., a large number of docking sites, instantaneous site repair and neurotransmitter clearance) it can be shown using LNA that

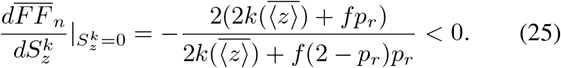

Thus, increasing 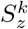 or including positive feedback will decrease *FF*_*n*_, and this decrease is stronger for large values of 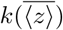 (left-hand plot in Fig 5). However, as can be seen from the right-hand plot in Fig 5, depending on the relative rates of the AP arrival frequency and neurotransmitter decay, negative feedback can reduce fluctuations in the level of neurotransmitters in the cleft. We confirm these results via exact stochastic simulation of the model, where for the parameter values considered, including negative feedback amplifies both *FF*_*n*_ and *FF*_*z*_, and positive feedback has the opposite noise buffering effect (Fig. 6).

**Fig. 6.**
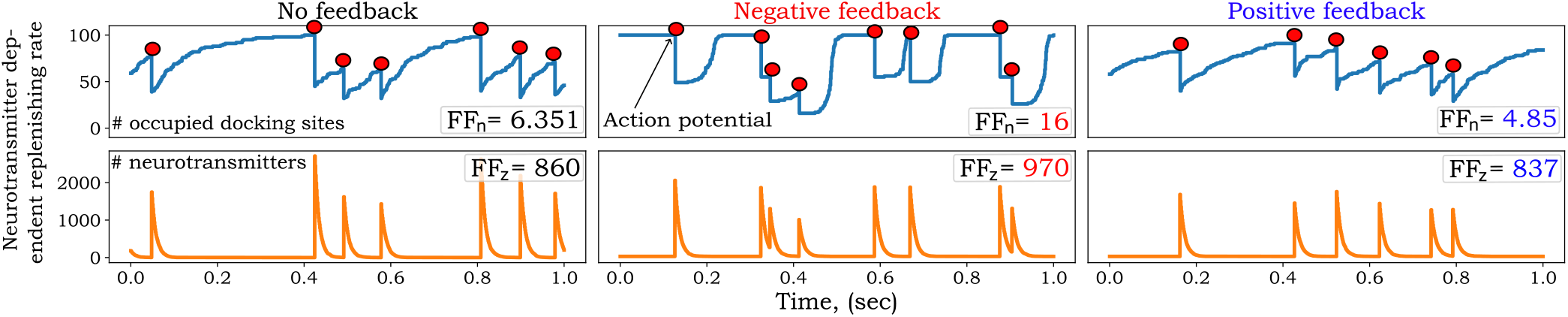
Neurotransmitter-dependent positive (negative) feedback on SV refilling rate reduces (increases) statistical fluctuation in neurotransmission. Three models are presented in this figure: the no feedback case with a constant refilling rate *k* = 10 sec^*−*1^ (left), a negative feedback model with a refilling rate that decreases with increasing neurotransmitter levels (middle), and a positive feedback model with a refilling rate that increases with increasing neurotransmitter levels (right). For the negative feedback, 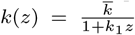, where 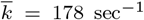, and *k*_1_ = 3. For positive feedback 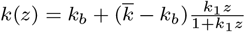, where 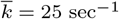, *k*_1_ = 0.001, and *k*_*b*_ = 7 sec^*−*1^ is a basal vesicle replenishing rate. These parameters were chosen such that across all feedback types the mean steady-levels are constant and approximately 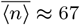 and 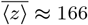. For each case, sample trajectories are shown and the Fano factor computed from a large number of simulations is shown on the respective plot. The red dots indicate a presynaptic AP and other parameters were taken as *f* = 10 *Hz, p*_*r*_ = 0.5, *M* = 100, *c* = 50 and *γ*_*z*_ = 100 sec^*−*1^. For this plot, we assume instantaneous docking site repair.

## VII. Conclusion

This contribution expands previous stochastic models of SV release in a chemical synapse on two different fronts: accounting for docking site repair after SV fusion; and inclusion of feedback regulation where released neurotransmitters can alter SV docking rates. In the former case, our results show that the repair process can be accounted for by defining an equivalent rate *k*_*eq*_ that is essentially the inverse of the average time for site repair and site re-loading by a SV. For a low number of docking sites, the steady-state mean and noise levels of docked SVs and released neurotransmitters only depend on the repair rate *k*_*r*_ via the equivalent rate *k*_*eq*_. In this case, varying *k*_*r*_ while keeping *k*_*eq*_ will not impact the steady-state statistical fluctuations. In contrast, for a large number of docking sites, the noise levels are amplified as a result of site repair for fixed *k*_*eq*_, and the amplification is the highest when *k*_*r*_ = 2*k*_*eq*_ (Fig. 4). However, our results show that these amplifications are quite modest, especially for the noise in the level of the neurotransmitter in the synaptic cleft. It is important to point out that while our model considers exponentially distributed repair times, future work can relax this assumption by considering non-exponential repair times.

We next consider feedback regulation in the model by considering that SVs dock at empty sites with a neurotransmitter-dependent rate *k*(*z*) – a monotonically increasing (decreasing) function *k* implementing positive (negative) feedback. Interestingly, both analytical small-noise approximations and stochastic simulations reveal that positive (negative) feedback will always attenuate (amplify) fluctuations in docked SV counts. To understand this effect intuitively consider the case when the arrival of the next AP is significantly delayed by random chance. Then in the absence of feedback (i.e., fixed SV docking rate), all docking sites will get occupied resulting in a large neurotransmitter release for the next AP. With positive feedback, the lowering of neurotransmitter levels due to the delayed AP will also decrease the docking rate resulting in a buffered neurotransmitter release. At the level of the neurotransmitter, our results show that depending on the parameter regime either feedback can provide effective noise attenuation, with positive feedback being more effective at low AP frequency, and negative feedback buffering fluctuations at high AP frequency (Fig. 5). Supported by these theoretical results, our work with experimental collaborators has characterized both positive and negative feedback loops involved in regulating the release of the dopamine neurotransmitter [38], [39]. Such feedbacks are implemented via auto-receptors on the neuron membrane that sense the neurotransmitter in the cleft and activate signaling pathways to modulate presynaptic parameters [40], [41]

Future work will consider a wider class of feedback where neurotransmitters can impact the release probability, the number of docking sites, or the time taken for site repair. Our preliminary work via stochastic simulations shows that negative feedback regulation of the release probability, i.e., the probability of release decreases in response to the buildup of neurotransmitters in the cleft can reduce both *FF*_*n*_ and *FF*_*z*_ (Fig. 7). On the mathematical front, nonlinearities introduced by such feedbacks in stochastic dynamical systems can be studied via different approaches, such as moment closure schemes [42], [43]. It will also be interesting to consider the time between successive APs that follow an arbitrary non-exponential distribution using the theory of renewal processes [44] with potentially time-dependent model parameters. This parametric time dependence captures a key physiological effect where a significantly delayed AP will lead to a lowering of SV docking rates and/or release probability due to reduced calcium levels in the axon terminal.

**Fig. 7.**
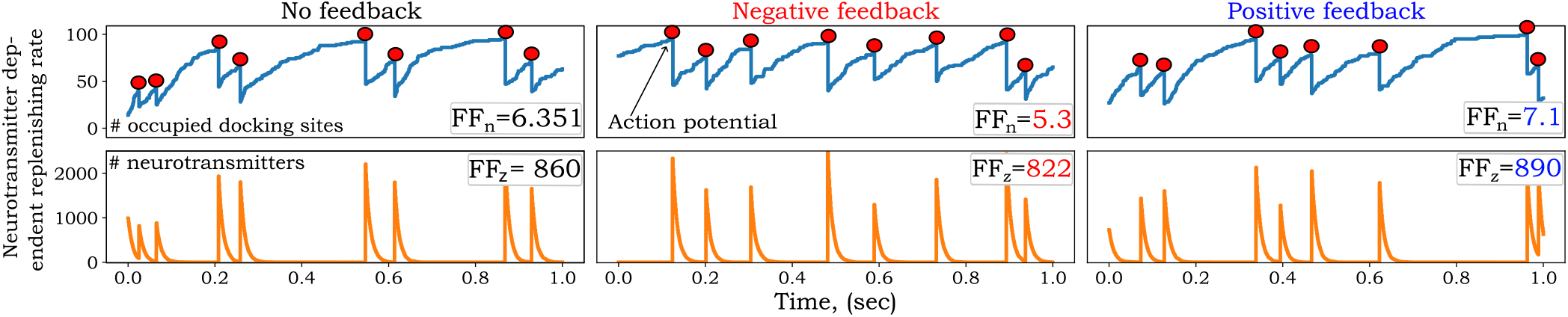
Neurotransmitter-dependent negative (positive) feedback on release probability reduces (increases) statistical fluctuation in neurotransmission. This figure is similar to Fig. 6 except that now the SV refilling rate is assumed to be constant *k* = 10 sec^*−*1^ and feedback is implemented on the probability of AP-triggered SV fusion. For the no feedback case *p*_*r*_ = 0.5 (left), for negative feedback (middle) 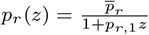, where 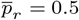 and *p*_*r*,1_ = 0.001. For positive feedback (right) 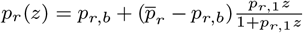, where 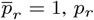, *p*_*r*,1_ = 0.001, and *p*_*r,b*_ = 0.5 is a basal vesicle release probability for positive feedback. In all cases, the mean steady-state levels were fixed at 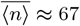 and 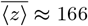. All other parameters were as in Fig. 6.

